# Collective dynamics of *Escherichia coli* growth under near-lethal acid stress

**DOI:** 10.1101/2024.10.17.618973

**Authors:** Rafael R. Segura Munoz, Victor Sourjik

**Affiliations:** Max Planck Institute for Terrestrial Microbiology, 35043 Marburg, Germany; Center for Synthetic Microbiology (SYNMIKRO), 35043 Marburg, Germany

**Keywords:** *Escherichia coli*, Enterobacteriaceae, acid resistance, collective behavior

## Abstract

Many neutralophilic bacteria, including *Escherichia coli*, can withstand acidic conditions due to the action of several protective mechanisms. While the survival of *E. coli* under growth-inhibitory extreme acid stress is well understood, less is known about the physiology of *E. coli* growth under severe but permissive acidity. Here, we observed that growth of *E. coli* MG1655 in a rich medium with pH of 4.1 to 4.4 exhibits a characteristic multi-phasic growth pattern, consisting of an initial exponential elongation in the absence of cell division, followed by growth arrest and subsequent resumption of exponential growth. The duration of the growth arrest phase was strongly dependent on the pH of the medium, but also on the initial cell density of the culture, suggesting the collective nature of this phenomenon. Cell-density dependent multiphasic growth at the near-lethal pH, including the initial increase in cell volume associated with either elongation or widening of the cell body, was also observed for all tested natural *E. coli* isolates and for *Salmonella enterica* serovar Typhimurium. Such transient increase in volume apparently enables bacteria to induce acid resistance systems, including the lysine-dependent Cad system, to subsequently modify pH of the medium in the density-dependent manner. Consistent with the collective recovery, even a minor fraction of acid-tolerant cells could fully cross-protect acid-sensitive cells enabling them to resume growth in coculture. Thus, collective dynamics plays a central role in bacterial growth under near-lethal acid stress.

**Importance:** Since many *E. coli* and other Enterobacteriaceae isolates are gastrointestinal pathogens for humans, it is important to understand their growth under acidic conditions imposed by the host physiology in the stomach, colon, and inside macrophages. Here we show that *E. coli* growth under near-lethal acidic conditions is a collective phenomenon that critically depends on cell density. This collective behavior is favored by changes in cell morphology as a response to high acidity. The observed density dependence might have implications for pathogen proliferation in the acidic environment of the human gastrointestinal tract and possibly also for their interactions with immune cells.

## Introduction

Neutralophilic bacteria grow optimally about two units below and above pH 7.0 (1). Some neutralophiles, such as *Escherichia coli*, have evolved multiple protective mechanisms that allow them to grow in acidic environments, such as the mammalian gastrointestinal tract and acidified water, soil, and food (2–8). The acid resistance of *E. coli* and other human pathogens has mostly been studied under growth-inhibiting, lethal acid stress (pH < 3.0) that simulates an infection-related passage through the stomach (9, 10), and to a much lesser extent under growth-permitting, near-lethal acid stress (pH 4.0-5.0) that mimics other common acidic environments (9, 11), such as the human duodenum, duodeno-jejunal junction and proximal colon (7, 8).

Among multiple protective mechanisms of *E. coli*, the most prominent are the amino acid-dependent acid resistance (AR) systems, which rely on H^+^ consumption during amino acid decarboxylation to raise the cytoplasmic pH (2–5). These include the glutamate-dependent Gad system (GDAR or AR2), the arginine-dependent Adi system (ADAR or AR3) and the lysine-dependent Cad system (LDAR or AR4) (2, 12). Each of these systems consists of a pH-dependent transcriptional regulator (GadE, AdiY or CadC, respectively), an amino acid decarboxylase (GadA/GadB, AdiA or CadA) and an antiporter (GadC, AdiC or CadB) that imports the respective external amino acid and exports the more alkaline product of amino acid decarboxylation, which is γ-aminobutyric acid (GABA) for the Gad system, agmatine for the Adi system, and cadaverine for the Cad system. These different systems are known to be induced under different conditions, with the Gad and Adi systems being particularly important under extreme acid stress, stationary phase, and/or anaerobic conditions, and the Cad system operating under mild (pH 5.0-5.8) and near-lethal acid stress (13). Moreover, a recent study demonstrated that operation of these systems may be partitioned within the same population of *E. coli* cells, with only fractions of cells expressing the Cad or Adi systems (13, 14). Such heterogeneity in the induction of stress response pathways indicates bet-hedging behavior, but possibly also division of labor and potential for collective behaviors in bacterial populations under acidic pH (13). Population density has been indeed shown to affect *E. coli* survival after exposure to lethal pH 2.0 (10, 15), but the nature of this effect remained unknown.

Here we show that, upon rapid exposing to near-lethal pH, strains of *E. coli* as well as *Salmonella enterica* serovar Typhimurium (*S. Typhimurium*) exhibits cell-density dependent multiphasic growth. In *E. coli* K-12, it consists of the initial phase of exponential elongation in the absence of cell division, followed by a transient cessation and a subsequent recovery of normal growth. The duration of this transient growth arrest was strongly dependent on cell density, apparently due to the collective deacidification of the medium that was partly dependent on the Cad system. The initial phase of the cell size increase was apparently important to enable the biosynthesis of the acid resistance systems, and it was also observed in other isolates of *E. coli* isolates and *S. Typhimurium*. Consistent with the collective deacidification, we observed that even a minor proportion of acid-tolerant *E. coli* cells is sufficient to support growth of the acid-sensitive *E. coli* population at near-lethal pH.

## Results

### *E. coli* exhibits density-dependent multiphasic growth under severe acid stress

To investigate the growth dynamics of *E. coli* at different values of acidic pH, we inoculated cultures at different inoculum sizes in LB medium with initial pH ranging from 7.0 to 4.0. We observed that the growth rate of *E. coli* K-12 strain MG1655 decreased progressively at lower pH (Fig. 1A-E and Fig. S1). Although the growth rate was apparently independent of the cell culture density at pH 7.0 or 5.0 (Fig. 1A, B), cell-density dependence of growth was observed starting at pH 4.4. Specifically, bacterial cultures exhibited multiphasic growth, including an initial, apparently exponential phase of increase in optical density (OD_600_) followed by an intermediate lag (mid-lag) phase and then a second exponential growth phase. At pH 4.4, such multiphasic growth was only observed for the lowest density culture (Fig. 1C), whereas high cell-density cultures showed continuous growth. At decreasing pH, the multiphasic growth became increasingly pronounced also for intermediate- and high-density cultures (Fig. 1D, E and Fig. S1A, B). No growth was observed at the pH 4.0 (Fig. S1C). Importantly, the duration of the growth arrest phase showed strongly pronounced dependence not only on pH but also on the cell density, with higher-density cultures exhibiting proportionally shorter or no lag phase.

**Fig. 1.**
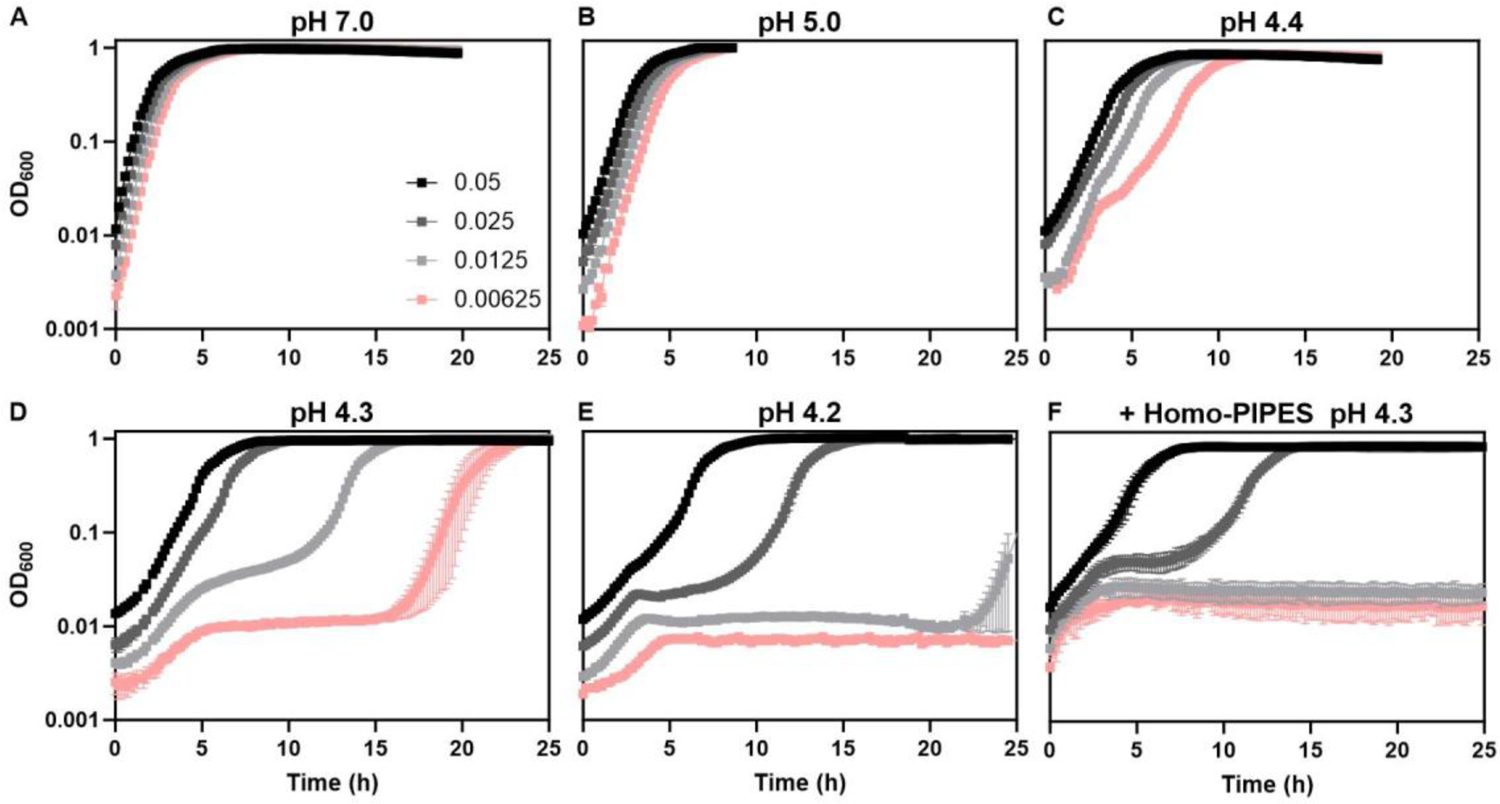
*E. coli* exhibits density-dependent growth at low pH. Growth of *E. coli* MG1655 measured using plate reader at indicated inoculum sizes in unbuffered LB adjusted to initial pH of (A) 7.0, (B) 5.0, (C) 4.4, (D) 4.3, (E) 4.2, or in (F) LB buffered with 33 mM Homo-PIPES at pH 4.3. Inoculum size values here and throughout correspond to OD_600_ measurements in a 10 mm cuvette using spectrophotometer, thus deviating from OD_600_ values measured in a plate reader. Mean and SEM of 3 replicates are shown.

We further investigated whether the density-dependent multiphasic growth is affected by the buffering capacity of the medium. Indeed, for *E. coli* cultures grown in LB with pH 4.3, buffered using homopiperazine-1,4-bis-2-ethanesulfonic acid (Homo-PIPES), the density dependence of growth shifted to higher-density cultures at intermediate buffering capacity (Fig. 1F), and growth entirely ceased at high buffering capacity (Fig. S2).

### Density-dependent growth is characterized by cell elongation and medium deacidification

Given the observed effects of buffering, medium deacidification may dictate the observed multiphasic growth. Consistently, the final pH of all cultures that reached high OD increased to >8.0, irrespective of their initial pH, while that of non-growing cultures remained low (Fig. 1 and Table S1). To elucidate how deacidification of the medium correlates with the multiphasic growth, OD and medium pH were monitored simultaneously for *E. coli* cultures with different inoculum size growing at an initial pH of 4.2 (Fig. 2 and Fig. S3). We observed that changes in pH were consistent with the cell growth (Fig. 2A-B and Fig. S3A-B). In particular, for low-density cultures where the multiphasic growth was pronounced, the exit from the intermediate lag phase coincided with the increase in pH. This resumption of cell growth occurred at pH ∼4.4, and it was preceded by a very slow gradual elevation of pH over the entire duration of the initial phase of OD increase and the mid-lag phase. In contrast, the high-density culture crossed this apparently critical pH value already during the first phase and thus did not enter the intermediate phase.

**Fig. 2.**
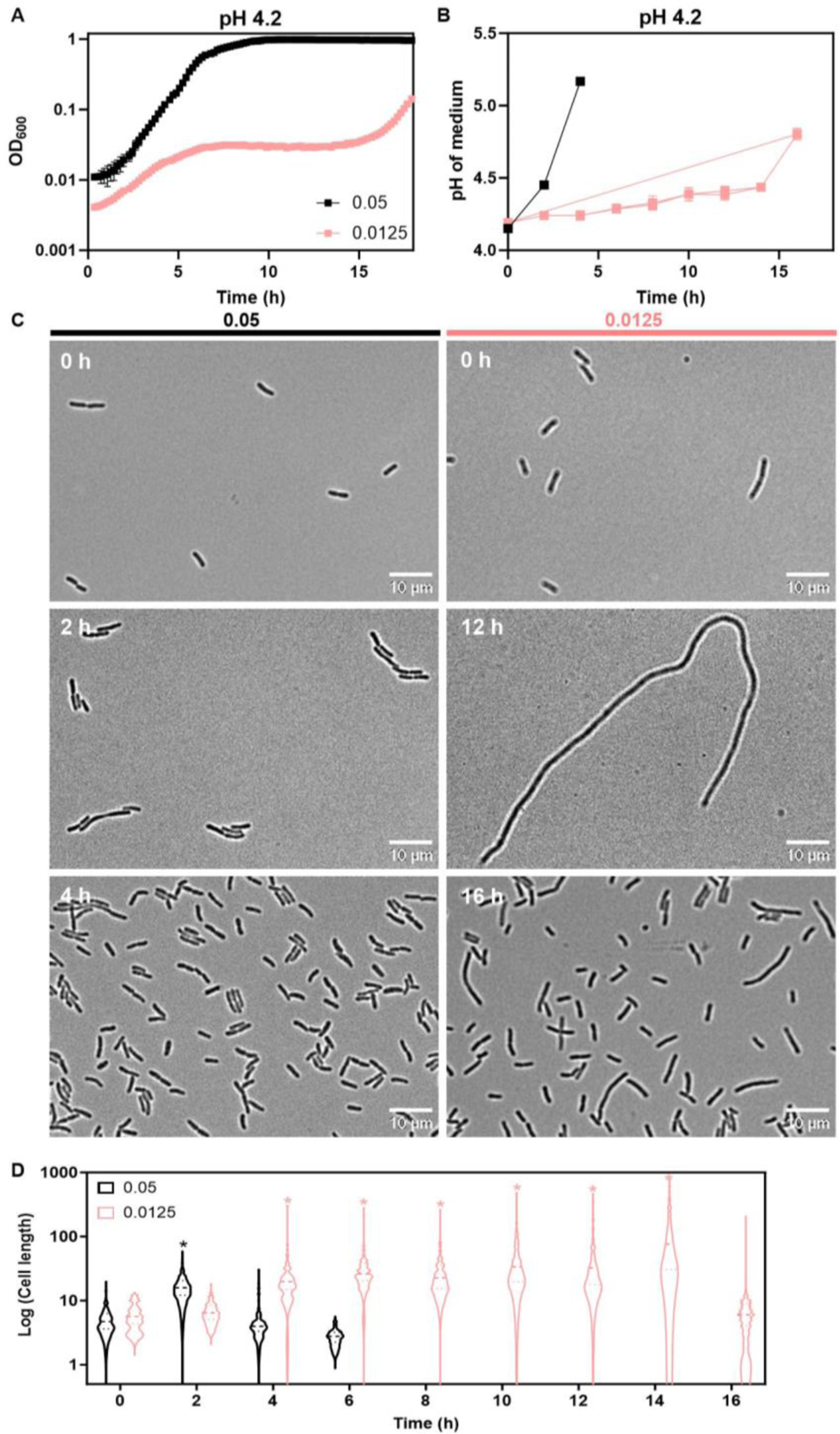
Multiphasic *E. coli* growth reflects changes of cell size and medium deacidification. *E. coli* MG1655 cultures with indicated inoculum size were grown in LB at pH 4.2. (A) Growth of the culture. (B) Medium pH, measured as described in Materials and Methods. Mean and SEM of 3 replicates are shown. (C) Morphology of *E. coli* cells at indicated time points. Scale bar is 10 µm. (D) Quantification of cell length for 100-150 cells at indicated time points, with the distribution of responses, the average (dashed line) and the upper and lower quartiles (dotted lines) being indicated. Significance of difference compared to *t* = 0, assessed using a paired *t*-test, is indicated by asterisks (\**p* ≤ 0.05).

To understand the nature of the initial phase of largely density-independent OD increase, we investigated changes in cell morphology during growth using microscopy. In all cultures growing at pH 4.2, cells showed pronounced but transient elongation, with its extent and the duration of the elongation phase dependent on the inoculum density (Fig. 2C, D and Fig. S3C). Cell length primarily increased during the first phase of growth, suggesting that this up to ∼10-fold increase in OD during the initial, apparently exponential growth phase, reflects the elongation of cells from the initial average cell length of 4.4 µm to ∼54 µm in the absence of cell division. The reduction of the cell length subsequently occurred concurrently with the resumption of growth after the mid-lag phase.

### Pre-induced non-growing cells are able to deacidify the medium

These results indicate that the density dependence of growth at pH below 4.4 could be explained by gradual, density-dependent deacidification of the medium over the net duration of the initial elongation and subsequent mid-lag phase, until the critical permissive pH above 4.4 is reached and the growth/division arrest is relieved. The ability of non-growing cells to deacidify the medium was further confirmed by monitoring pH in a culture of translation-inhibited cells (Fig. 3). Here we compared *E. coli* cultures that were pre-grown in LB at either neutral or acidic (4.2) pH for 1 hour and subsequently transferred to LB containing chloramphenicol at pH 4.2 for 3 hours to inhibit translation. The density of both cultures was adjusted to OD_600_ = 0.1 to reflect the cell density at the mid-lag phase of the 0.0125 inoculum culture (Fig.2A), and OD_600_ and medium pH were monitored. We observed that translation-inhibited cells pre-incubated in acidic medium did not grow but increased medium pH to approximately 4.4 in 14 hours (Fig. 3A-B). This rate of deacidificaiton was similar to that observed during the mid-lag phase of growing cultures (Fig. 2B and Fig. 3C). In contrast, the cultures that were not pre-exposed to acidic pH prior to the inhibition of translation could not efficiently change pH of the medium, suggesting that the growth-dependent pre-induction of the AR system(s) may be essential for subsequent deacidification of the medium by non-growing cells.

**Fig. 3.**
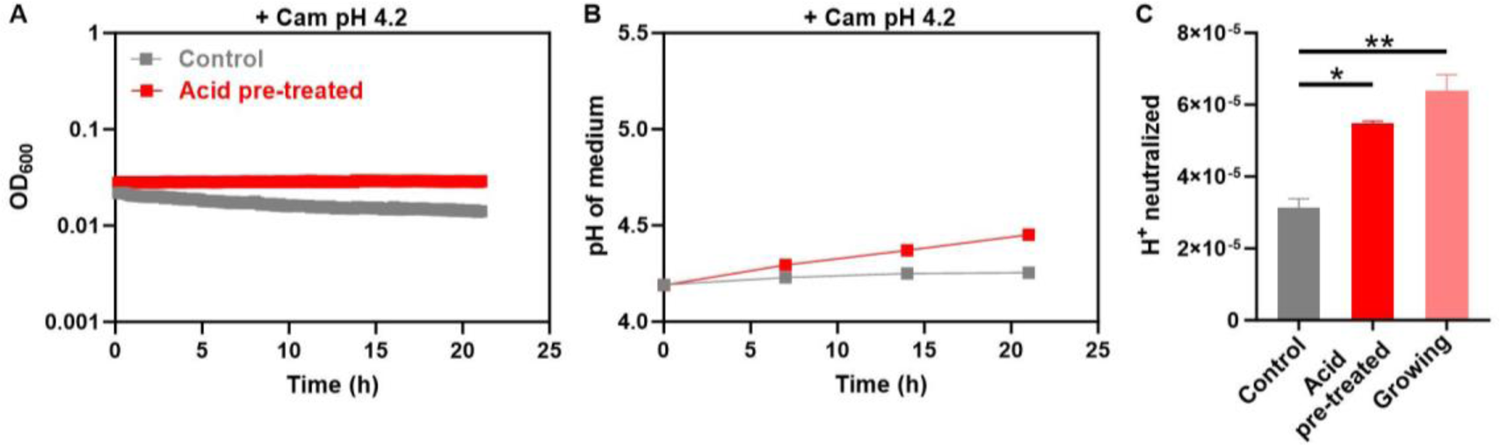
Non-growing cells can efficiently deacidify the medium. *E. coli* cells treated with chloramphenicol to inhibit translation were incubated in LB at pH 4.2. (A) Growth of the culture. (B) Medium pH. Inoculum size was 0.1. Prior to the chloramphenicol (Cam) treatment, cells were exposed to either pH 4.2 (acid pre-treated) or pH 7.0 (control) for 1 h in LB. (C) Rate of neutralization, calculated as the molar mount of protons neutralized per OD_600_ per hour in translation-inhibited cells, or in untreated cells during the mid-lag phase (data from Fig. 2B). Mean and SEM of 3 replicates are shown. Significance of differences, assessed using an unpaired *t*-test, is indicated by asterisks (\**p* ≤ 0.05, \*\**p* ≤ 0.01).

### Importance of individual AR systems in multiphasic growth under acid stress

To assess which AR system(s) contribute to the acid tolerance of *E. coli* during multiphasic growth under near-lethal acid stress, we first tested *E. coli* strains deficient in the expression of individual AR systems. A strain lacking *cadC*, encoding the transcriptional activator of the AR4 system, showed much reduced growth in LB pH 4.3 even at high inoculum size (Fig. 4A), suggesting reduced acid tolerance of this mutant. In contrast, no effect was observed for strains deleted for genes encoding the transcriptional activators of AR2 (Δ*gadE*) or AR3 (Δ*adiY*). Consistent with this, supplementation of LB pH 4.2 with lysine, the substrate of AR4, resulted in a strong improvement in growth (Fig. 4B and Fig. S4), with a reduced duration of the mid-lag phase. The duration of the mid-lag phase was also shortened in *E. coli* cells overexpressing *cadC* (Fig. 4C). Although the addition of glutamate (substrate of AR2) even caused a growth reduction (Fig. S4), the overexpression of *gadE* could improve growth at low pH (Fig. S5).

**Fig. 4.**
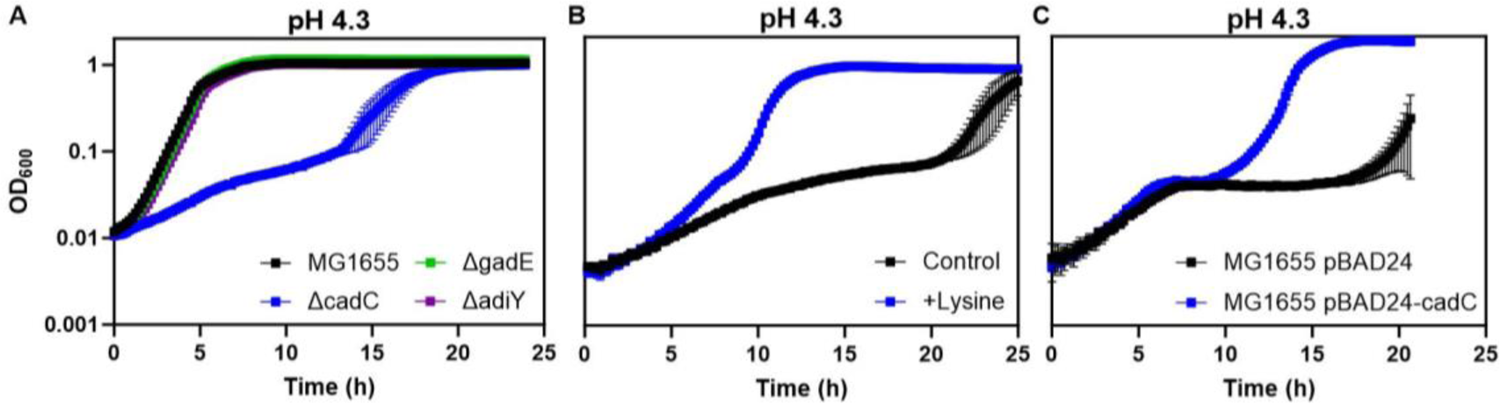
Cad system is involved in cell density-dependent acid tolerance. (A) Growth of *E. coli* MG1655 wild type or *ΔcadC*, *ΔgadE*, or *ΔadiY* mutants, as indicated, in LB at pH 4.3. Inoculum size was 0.05 for all strains. (B) Growth of *E. coli* MG1655 in LB with or without 20mM lysine supplementation, at inoculum size of 0.0125. (C) Growth of *E. coli* MG1655 carrying either pBAD24 or pBAD24-*cadC*, at inoculum size of 0.0125. Mean and SEM of 3 replicates are shown.

### Other *E. coli* and *Salmonella* strains exhibit cell-density dependent growth and morphological changes at near-lethal pH

To test whether other *E. coli* isolates also exhibit cell-density dependent multiphasic growth at near-lethal pH, we monitored the growth dynamics of two uropathogenic *E. coli* strains, UTI89 (16) and CFT073(17) (Fig. 5 and Fig. S6). Although these strains showed slightly higher acid tolerance than MG1655, the dependence of growth on cell density was observed for both strains at pH of 4.1 (Fig. 5A, B and Fig. S6A, B). However, the initial phase of the multiphasic growth was less pronounced, and in contrast to MG1655 these strains primarily exhibited biomass accumulation in the center of the cells and only minor elongation (Fig. 5C-H). Nevertheless, also for these strains the resumption of the normal growth coincided with the restoration of the normal cell size and with increase in pH (Fig. 5E-H and Fig. S6C, D; high-density culture).

**Fig. 5.**
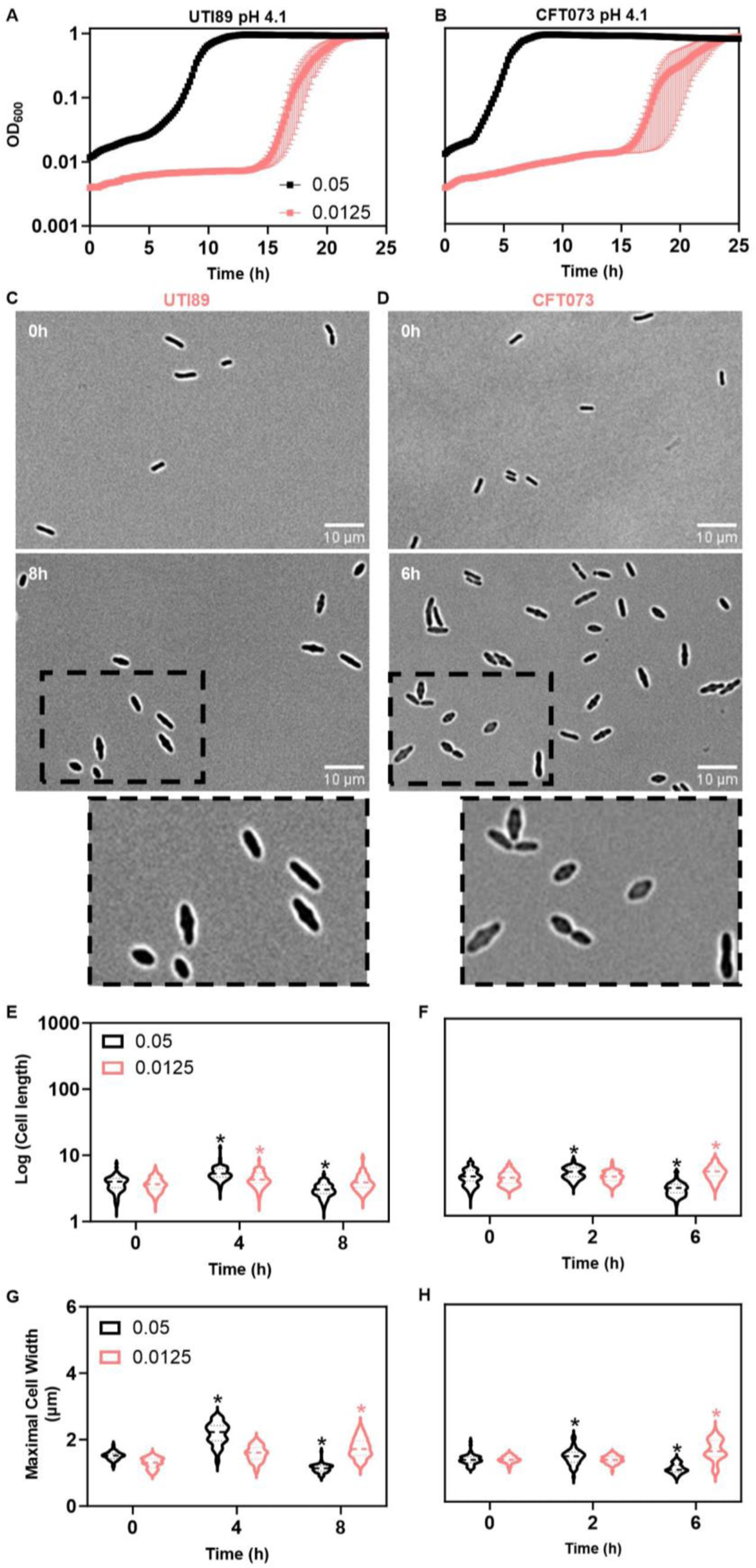
Uropathogenic *E. coli* strains exhibit cell-density dependent growth and morphology changes at near-lethal pH. Growth of *E. coli* strains UTI89 (A) and CFT073 (B) at indicated inoculum sizes in LB at pH 4.1. Mean and SEM of 3 replicates are shown. Morphology of UTI89 (C) and CFT073 (D) at indicated time points. Scale bar is 10 µm. Dotted lines specified the magnified area. Quantification of cell length for 100-150 cells of UTI89 (E) and CFT073 (F), and of maximal cell width for UTI89 (G) and CFT073 (H), done and represented as in Figure 2.

Given this difference in the acid-induced phenotype between the K-12 and UTI89 and CFT073 strains, we further tested morphological changes for a number of strains from the ECOR collection of natural isolates of *E. coli* (18) grown in LB at pH 4.1. Both cell elongation and widening, or their combination, were observed in the ECOR strains belonging to phylogroups A (which also include *E. coli* K-12), B1 and D (Fig. S7A, B and D). Strains from B2 phylogroup showed primarily cell widening (Fig. S7C), consistent with the phenotype of UTI89 and CFT073 that also belong to this group.

Because *Salmonella* and *E. coli* species share acid resistance mechanisms (2, 5, 19), we further tested growth of *S. Typhimurium* at conditions ranging from neutral to near-lethal acidic pH. We observed the growth rate reduction with lower initial medium pH, although at lower pH than for *E. coli* strains, as well as the density-dependence of growth (Fig. S8 and Fig. S9A). The initial growth phase was less pronounced in *S. Typhimurium* than in *E. coli* K-12, but there was both a transient change in the cell size (Fig. S9C-D) and an increase in the medium pH (Fig. S9B) before resumption of normal division.

### Cross-protection between acid-tolerant and acid-sensitive strain in a mixed population

Given our observation that *E. coli* growth at near-lethal pH relies on collective deacidification of the medium, we hypothesized that, in a coculture, acid-sensitive cells could be complemented for growth by acid-tolerant cells. To test for such cross-complementation, we cocultured wild-type (WT) and Δ*cadC E. coli* MG1655 at equal amounts (WT 50: KO 50) in LB at pH 4.3, and compared growth of this coculture to the monocultures of either WT or Δ*cadC* (Fig. 6). Consistent with our previous results (Fig. 4A), growth of the Δ*cadC* monoculture at low pH was severely reduced (Fig. 6A) and no deacidification of the medium was observed over 9 h of incubation (Fig. 6B). Overall growth and medium deacidification for the WT 50: KO 50 coculture was intermediate between the WT 100 (i.e., twice the WT density) and WT 50 monocultures, suggesting that Δ*cadC* cells contribute to medium deacidification, probably via other AR systems. Most importantly, quantification of the colony-forming units (CFUs) revealed nearly identical growth of the WT and Δ*cadC* strains in the coculture at each time point, further confirming the entirely collective nature of the acid tolerance under these conditions. Consistently, growth complementation was also observed in the coculture of WT 20: KO 80 (Fig. S10), demonstrating that even a minor fraction of acid-tolerant bacteria can support the growth of the entire population at low pH.

**Fig. 6.**
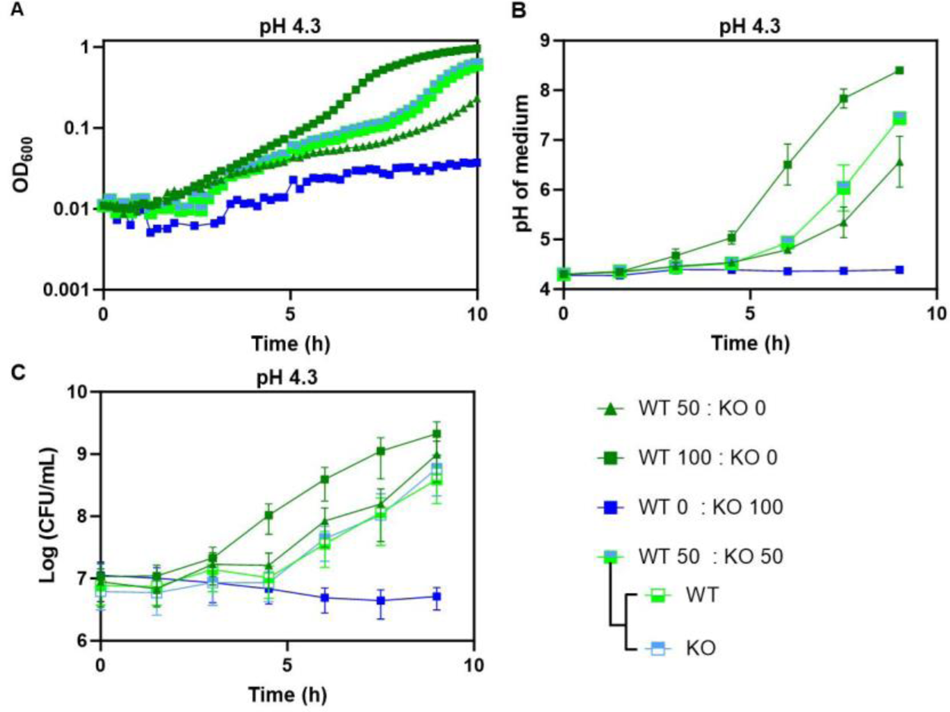
Acid-tolerant strain complements growth of an acid-sensitive strain in a coculture. Acid-tolerant (WT; MG1655 carrying pBAD24) and sensitive (KO; MG1655 *ΔcadC*) were cocultured at total inoculum size of 0.05 in LB at pH 4.3. Two strains were mixed at equal proportions (50:50); monocultures were inoculated either at half the density or at the full density. (A) Growth of the cocultures. (B) Medium pH. (C) Colony-forming units (CFU) per mL or indicated cultures. The CFUs for WT and KO were distinguished by plating cocultures on LB agar with ampicillin and kanamycin, respectively. Mean and SEM of 3 replicates are shown.

## Discussion

Many neutralophilic bacteria, including pathogens, are naturally exposed to acidic environments such as the mammalian colon and acidified foods, soil and water (2–6). To date, *E. coli* growth at near-lethal pH has only been studied for growth or no-growth outcomes (15) or for the overall reduction in growth rate (4, 11, 12, 15). Here, we systematically examined the dependence of *E. coli* growth not only on pH but also on cell density.

We observed that *E. coli* growth at the near-lethal acid pH is multiphasic, consisting of an initial phase of optical density increase, a mid-lag phase of apparent growth arrest, and a phase of growth resumption. This multiphasic growth was not solely determined by the medium pH, but also by the cell density of the culture. While the first and the third phases of growth showed little density dependence, low cell density cultures displayed higher duration of the mid-lag phase compared to high cell density cultures. Our results suggest that this density dependence could be explained by gradual collective deacidification of the medium during the mid-lag phase, which allows the resumption of growth above pH of 4.4. This deacidification depends on the number of deacidifying cells, but also on the buffering capacity of the medium, and on the presence of the AR systems, primarily AR4 (Cad) system.

We could further show that the ability of *E. coli* cells to deacidify the medium depends on their pre-exposure to acidic pH, likely because of the need to induce the expression of acid resistance system(s). Consistent with that, the non-growing, translation-inhibited *E. coli* cells could deacidify the medium similarly to the deacidification observed during the mid-lag phase of growth, but only when pre-exposed to acidic pH before the inhibition of translation. We propose that, during the multiphasic growth of the culture, this induction of the acid resistance systems occurs during the initial, density-independent phase of increase in the optical density. In all tested strains of *E. coli* as well as in *S. Typhimurium*, this first stage corresponded to changes in cell morphology. For *E. coli* MG1655 where it was most pronounced, we observed an extensive (up to ∼10 fold) cell elongation that could account for a similar-fold increase in optical density. Such elongation at low pH is consistent with a recent report that described it for several *E. coli* strains and for *Citrobacter rodentium* but not for *Salmonella*, and explained it by the inhibition of the essential divisome protein FtsI (penicillin-binding protein 3, PBP3) that is involved in septal peptidoglycan biosynthesis (20). Our observation that some *E. coli* strains, particularly those from B2 phylogroup that contains many uropathogens, increase in the cell width suggest that in those instances lateral rather than septal biosynthesis of peptidoglycan is inhibited at low pH. Nevertheless, in all cases, also including *Salmonella*, an increase in the cell size correlating with the initial phase of optical density increase is observed. Our results suggest that this increase of the biomass may be required to allocate additional protein resources in acid resistance. Notably*, E. coli* and *Salmonella* are known to transiently increase cell size in response to other stress conditions (21–23), indicating that the morphological response described here may be a general mechanism to enable bacteria to allocate resources in stress response.

The observed collective acid tolerance of *E. coli* growth at near-lethal pH may be related to the recently shown heterogeneous expression of the Cad system due to the low copy number of its regulator CadC (13, 24). It was suggested that stochastic differentiation of *E. coli* population into distinct *cad-on* and *cad-off* subpopulations (25) might represent division of labor in a bacterial population (13). Our results indeed support this hypothesis, demonstrating that even a minor fraction of Cad-positive cells is sufficient to enable growth of the entire population.

Taken together, here we could identify collective dynamics and morphological changes that are important for *E. coli* growth under acidic conditions, contributing to understanding of the mechanisms that enable *E. coli* strains to invade acidic intestinal microenvironments (26–28). The observed response could also affect bacterial interactions with immune cells, by counteracting phagocytosis (29) and/or by enabling survival within the acidic phagosome (26, 30).

## Materials and methods

### Bacterial strains and plasmids

All strains and plasmids used in this study are listed in Table S2. Genetically modified strains were derived from *E. coli* MG1655 using P1 transduction (31). pTrc99a-gadE was constructed by PCR amplification of *gadE* from MG1655 genome followed by Gibson assembly to clone it into pTrc99a expression vector inducible by isopropyl β-d-1-thiogalactopyranoside (IPTG).

### Culture conditions

Bacteria were cultured in lysogeny broth (LB) medium (10 g/L tryptone, 5 g/L yeast extract, and 5 g/L NaCl) at pH 7.0 (unless otherwise stated). HCl was used to adjust pH. Where indicated, lysine or other amino acids were added to a final concentration of 20 mM before adjusting the pH to 4.3. Pre-cultures were prepared by inoculating 2% overnight culture into LB and incubated for ∼1.5 hours until OD_600_ 0.50 - 0.70 at 200rpm and 37 °C. Cultures were washed three times (1500 x *g*, 3 min, 20 °C) using medium with the target pH and OD_600_ was adjusted to 0.50 using a 10 mm cuvette (REF 67742, Sarstedt). These OD-adjusted cultures were inoculated into LB with the target pH at 10% (OD_600_=0.05), 5% (OD_600_=0.025), 2.5% (OD_600_=0.0125), or 1.25% (OD_600_=0.00625) to generate inoculum sizes of ∼1.00×10^7^ cells/mL, 5.00×10^6^ cells/mL, 2.50×10^6^ cells/mL and 1.25×10^6^ cells/mL, respectively. HCl was utilized to acidify the medium. Growth was monitored in 48-well plates (500 µL/well, REF 677102, Greiner bio-one) in a plate reader (Infinite, Tecan GmbH) using alternating cycles of 150 s linear and orbital shaking at 37°C. When indicated, homopiperazine-1,4-bis(2-ethanesulfonic acid) (AB117062, Homo-PIPES, abcr GmbH) was added as buffer.

For pH measurements, the lids of 48-well plates were pierced with a sterile twisted drill bit to make sampling holes in each of the wells. The border of the plate and the holes in the wells were then sealed with parafilm to prevent evaporation. Samples were taken at specified intervals and pH was measured using a standard pH meter (ThermoScientific) or an Ultra-Micro-ISM pH meter (Mettler-Toledo) for small volumes. Quantification of CFUs for coculture experiments was done by plating serial dilutions on LB agar plates containing the appropriate antibiotics and manual counting of colonies.

### Test of pH changes by translation-inhibited cells

Pre-cultures grown as above were first transferred to LB medium with either pH 4.2 or pH 7.0 and incubated at 200 rpm for 1 hour at 37°C. Cells were then transferred to LB containing 200 mg/mL chloramphenicol (Cm) at pH 4.2 and incubated at 200 rpm, 37 °C for 3 hours. The OD_600_ was then adjusted to 0.50, the cultures were inoculated at the specified cell density and OD_600_ and pH were monitored during incubation in the plate reader as described above.

### Bright field microscopy

Cells were collected from 50 µL of culture by centrifugation (4500 *g*, 1 min, 20 °C) and the supernatant was used for pH measurement. The pellet was immediately resuspended in 10 µL PBS pH 5.0. A 5µL aliquot was placed on an agarose pad for microscopy.

Images were captured with a 20X objective using Zeiss observer Z1 or a Zeiss Elyra 7. Length and width of 100 to 150 cells were measured using Fiji 1.53 segmented lines tool.

### Growth complementation assay

Cultures adjusted to OD_600_ = 0.50 were used to generate 50:50 and 20:80 cocultures of acid-tolerant and resistant strains at a total density of OD_600_ = 0.05 in LB pH 4.3. At each time point, serial dilutions were made and colonies were counted on LB agar plates containing the appropriate antibiotics.

## Acknowledgment

We thank Kristin Jung and Sophie Brameyer for providing pBAD24-cadC and for discussions, and Rudolf K. Thauer for discussions. R.R.S.M. was supported by a postdoctoral fellowship from the Alexander von Humboldt (AvH) Foundation.

## Author contributions

Rafael R. Segura Munoz, Conceptualization, Formal analysis, Funding acquisition, Investigation, Methodology, Visualization, Writing – original draft, Writing – review and editing | Victor Sourjik, Conceptualization, Funding acquisition, Project administration, Supervision, Writing – original draft, Writing – review and editing.

## Data availability

All of the data are included in this article.

## Abbreviations

GDAR: glutamate-dependent Gad system

ADAR: arginine-dependent Adi system

LDAR: lysine-dependent Cad system

Homo-PIPES: homopiperazine-1,4-bis(2-ethanesulfonic acid)

Cm: Chloramphenicol.

